# DNA: A Nanoscale Archimedes’ Screw

**DOI:** 10.64898/2026.06.25.734429

**Authors:** Xavier Mleziva, Christopher Maffeo, Aleksei Aksimentiev

## Abstract

Rotating helices have been utilized for many purposes, including the transport of solid material and fluids within man-made machines, for a little over two millennia. Here, we show that the rotation of a biological helical molecule—a DNA duplex—can move water and ions through a nanoscale pore. While the rotation-induced flow of water is generated by the steric shape of the DNA molecule, an even faster transport of cations is caused by electrostatic interactions. The rotation-induced ion flux is found to depend on the cation type, offering potential utility for ion separation. Finally, we show that the torque-driven duplex can move ions against a concentration gradient, realizing the Archimedes screw principle at the nanoscale.

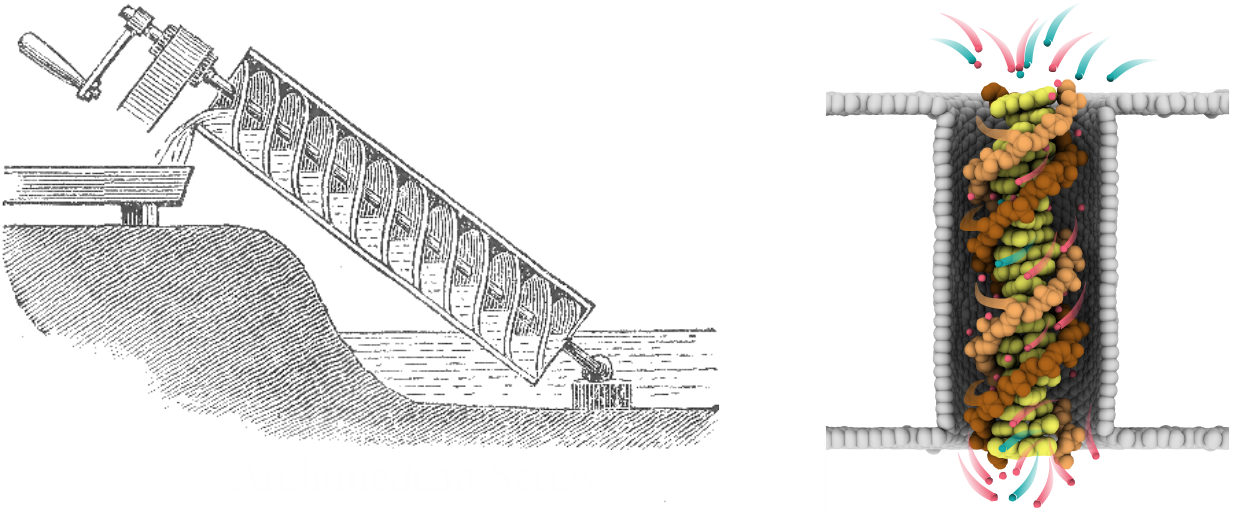

---

Throughout history, pumps have been essential components of human technology, performing tasks that range from moving water, waste, or even medicine^1^ to heating buildings. On the macroscale, fluid flow is described by Bernoulli’s law,^2^ which macroscopic pumps modulate by either a mechanical change of the volume occupied by the fluid, as in a syringe, or by imparting momentum on the fluid, as in a swimming pool pump.^3^ And while the efficiency of macroscale pumps has been optimized by centuries of development, scaling down macroscopic pumps to nanometer dimensions presents considerable challenges.^4^ The pitfalls of macroscale pump miniaturization stem from the fact that the physical forces dominating macroscopic fluid motion, such as gravity and inertia, are negligible at the nanoscale.^5^

The promise of lab-on-chip technologies^6^ has stimulated the development of microfluidic pumps using conventional microfabrication techniques, such as deep reactive ion etching, metal deposition, and resin printing.^7^ The fabricated micropumps generate microfluidic flow via a variety of mechanisms that may rely on passive processes, such as osmosis or surface tension effects, or require external energy input in the form of heat, electromagnetic field, or pressure gradient.^7^ Further miniaturization of pumps has become possible using nanopores of various geometries ^8,9^ and surface properties.^10^ Nanopumps of this type were used to generate molecular flow driven by light and pH,^9^ electro-osmosis^8,11^ and photogenerated electric fields.^12^ While not being able to move fluids, the smallest man-made pumps are chemically synthesized macromolecules that can directionally move individual chemical moieties using catalytic^13^ or ratchet mechanisms^14^ alongside photochemical,^15–17^ transamidation,^18^ or redox reactions.^19,20^

Within the realm of biology, evolution has transformed polypeptide chains into highly efficient molecular pumps that transport ions,^21,22^ protons,^23^ and other molecular species^24^ across cellular membranes. Such proteins use the energy of ATP hydrolysis to cyclically modulate their structure, opening and closing transmembrane passages while capturing and releasing specific molecules at their destination sides of a cellular membrane. One particular class of biological pumps, the V_0_V_1_ ATPases, utilizes mechanical rotation of the V_0_ domain relative to the V_1_ domains to pump protons^25,26^ and ions^27^ across the membrane and against the concentration gradient. The action of such a motor superficially resembles that of a macroscale pump but involves a dramatically different operating principle—the molecular ratchet.^28,29^ These rotary molecular motors can also operate in reverse. Thus, F_0_F_1_ ATP-synthase can use a transmembrane proton concentration gradient to rotate its transmembrane F_0_ domain relative to the solvent exposed F_1_, mechanically driving ATP synthesis.^30,31^ As such rotation can also be driven by a transmembrane electric potential gradient of electric potential,^32^ the operation of F_0_F_1_ ATP synthase resembles that of a macroscale electromotor.

All-atom molecular dynamics (MD) simulations have become a convenient tool to probe and design nanoscale molecular pumps.^33–35^ Thus, MD simulations of fixed-geometry nanotubes showed that molecular flux can be produced by thermal gradients,^36,37^ electro-osmosis,^38,39^ and exothermic reactions.^40^ Unidirectional water pumping was computationally realized via mechanical modulation of a nanotube’s shape.^41^ An electrically neutral screw-shaped nanochannel was shown to be capable of pumping water,^42^ whereas an engineered version of a biological water channel— aquaporin—of pumping ions in alternating electric field.^43^ In parallel, the robust programmability of DNA hybridization interactions have made DNA a material of choice for building complex nanoscale structures and mechanisms.^44^ Specifically, the DNA origami method^45^ was used to build complex rotary apparatuses^46–48^ and use external electric field or water flow to power unidirectional rotary motion.^49,50^ At the same time, MD simulations have shown that a single DNA duplex can act as a capable electromotor,^51^ a computational prediction confirmed by nanopore translocation experiments.^52^

Here, we use the all-atom MD method to investigate to what extent the action of a DNA duplex electromotor is reversible, that is, if a rotating DNA helix can pump water or ions, similar to its biological brethren F_0_F_1_ ATP synthase. For this purpose, we built an all-atom model of a 21basepair (bp) DNA duplex, placed it inside a carbon nanopore and solvated the system with 1 M KCl solution, Fig. 1a. The application of a 50 pN nm torque to phosphorus atoms of the DNA about its axis produced DNA rotation at a rate of 0.375 rev/ns, Fig. 1b and Supporting Video S1. The rotation rate was found to scale linearly with the applied torque, Fig. 1b, in accordance with the previous work.^51^ In the absence of applied torque, the DNA duplex was seen to undergo stochastic rotations driven by thermal fluctuations, Supporting Video S2, whereas reversing the sign of the torque reversed the DNA rotation direction Supporting Video S3.

**Figure 1:**
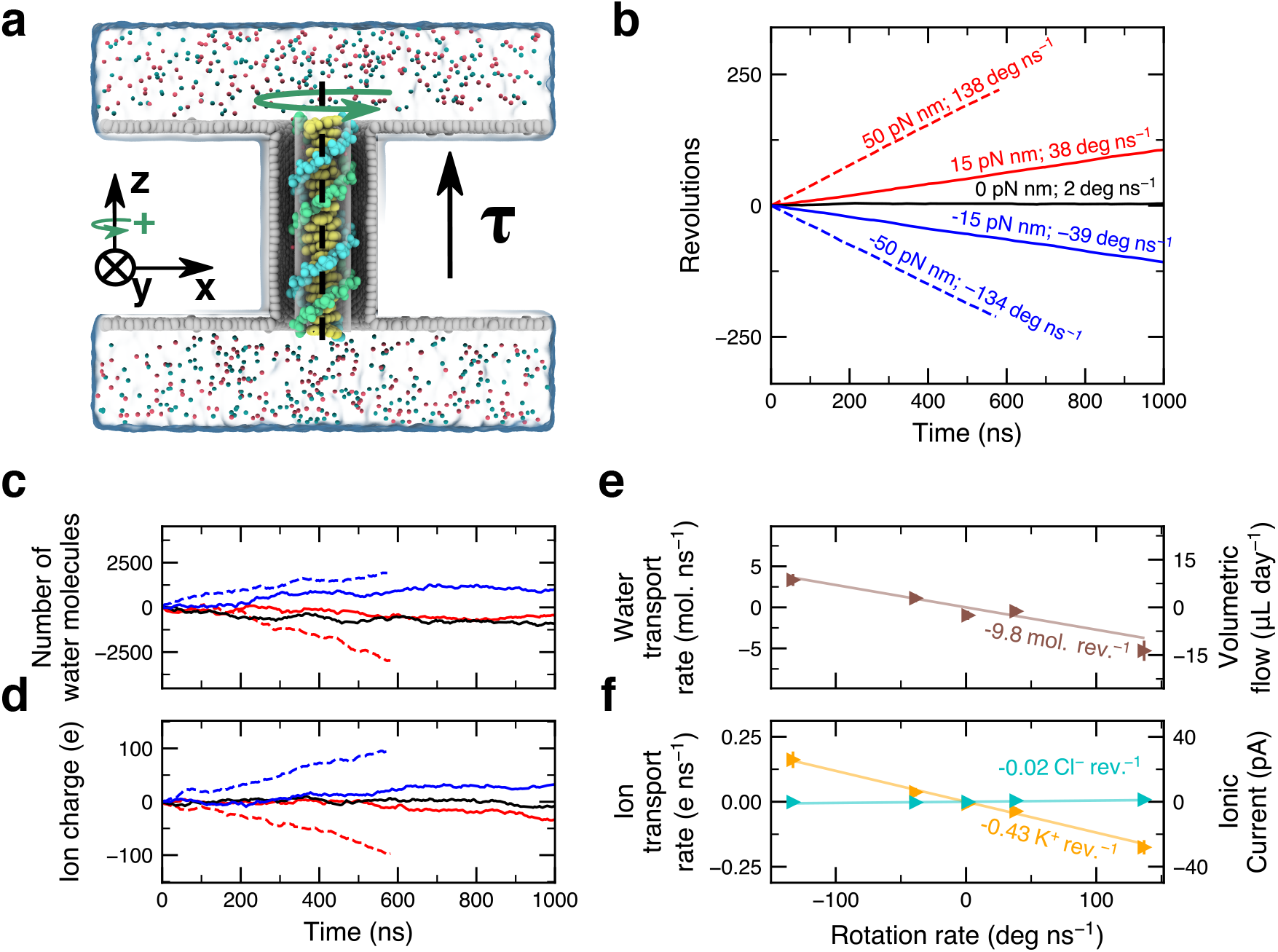
DNA rotation produces water and ion flux through a nanopore. **a**, Simulation system containing a 21 basepair DNA helix (yellow for bases; light blue and green for backbone) placed within a carbon nanotube 6.5 nm in length and 3.3 nm in diameter, submerged in 1 M KCl electrolyte. The nanopore surface is electrically neutral. Torque *τ* is applied along the nanopore axis (*z* axis) to the phosphorus atoms of the DNA duplex, producing rotation of the DNA duplex. The phosphorus atoms of the DNA are restrained to move along the surface of a cylinder coaxial with the nanopore (see SI Methods for details). **b**, Angular displacement of the DNA duplex *versus* simulation time at several values of applied torque. The value of the applied torque and the average angular velocity of the duplex (the slope of each curve) are indicated in matching colors. **c, d** Number of water molecules (panel c) and the total charge of ions (panel d) transported through the nanopore over the course of MD simulations carried out at different values of applied torque. The color and the pattern of the lines correspond to the applied torque as specified in panel b. The positive direction of transport is defined in panel a. Panels b-d, depict 4 ns block averaged data from MD trajectories each lasting at least 500 ns. **e**, Water transport rate (left axis) and the volumetric water flow (right axis) as a function of DNA rotational velocity. **f**, Ion transport rate (left axis) and the ionic current (right axis) as a function of DNA rotational velocity. In panels e and f, each data point represents an average over an MD trajectory of at least 500 ns. The error bars represent the standard error of mean calculated after splitting data into 50 ns blocks.

The forced rotation of the DNA duplex was found to generate a flow of water and ions through the nanopore in the direction opposite to that of the applied torque, Fig. 1c,d. The rate of water transport was found to scale linearly with the DNA rotation velocity, and hence with the applied torque. Approximately 9.8 water molecules were transported per revolution of the helix, Fig. 1e, regardless of the rotation rate. For the nanopore dimensions considered in the simulation, fully evacuating the nanopore volume occupied by water requires approximately 102 revolutions of the helix.

In addition to transporting water, rotation of the DNA duplex produced a directional transport of ions. Approximately 0.43 K^+^ ions were transported per revolution of the helix in the direction opposite to that of the applied torque, whereas the rate of Cl^*−*^ transport was close to zero, Fig. 1f. In comparison to the transport of water, the lower molecular flux of cations is explained by the 30-fold lower number density of the latter, whereas the nearly zero flux of anion is caused by the negative charge of the DNA duplex, which prevents anions from entering the nanopore. Consequently, the ionic current generated by the rotation of the duplex is dominated by cations, with the current magnitude scaling linearly with the applied torque, Fig. 1f. For the nanopore system considered in our simulations, a torque of 50 pN nm (similar to that generated by an ATP synthase^53^) generates an ionic current of a 30 pA magnitude, which is measurable using current electrophysiology instruments.^54^

To determine the molecular mechanism responsible for the generation of water and ion flow, we constructed an all-atom model of a 21 bp DNA duplex fully confined within a 3.3 nm diameter nanotube. Both the DNA duplex and the nanotube were coaxially aligned with one another and covalently bonded to themselves over the periodic boundary of the system, Fig. 2a. The number of water and ions placed within such an effectively infinite nanopore system was guided by the analysis of the equilibrated finite-size system, Fig. 1a. Upon brief equilibration, external torque was applied to the DNA duplex to produce its rotation. The radial profile of water density did not exhibit a dependence on the rotation direction, Fig. 2b, and matched the previously described distributions,^55^ including the increased water density near the nanotube surface caused by water adsorption.^56,57^

**Figure 2:**
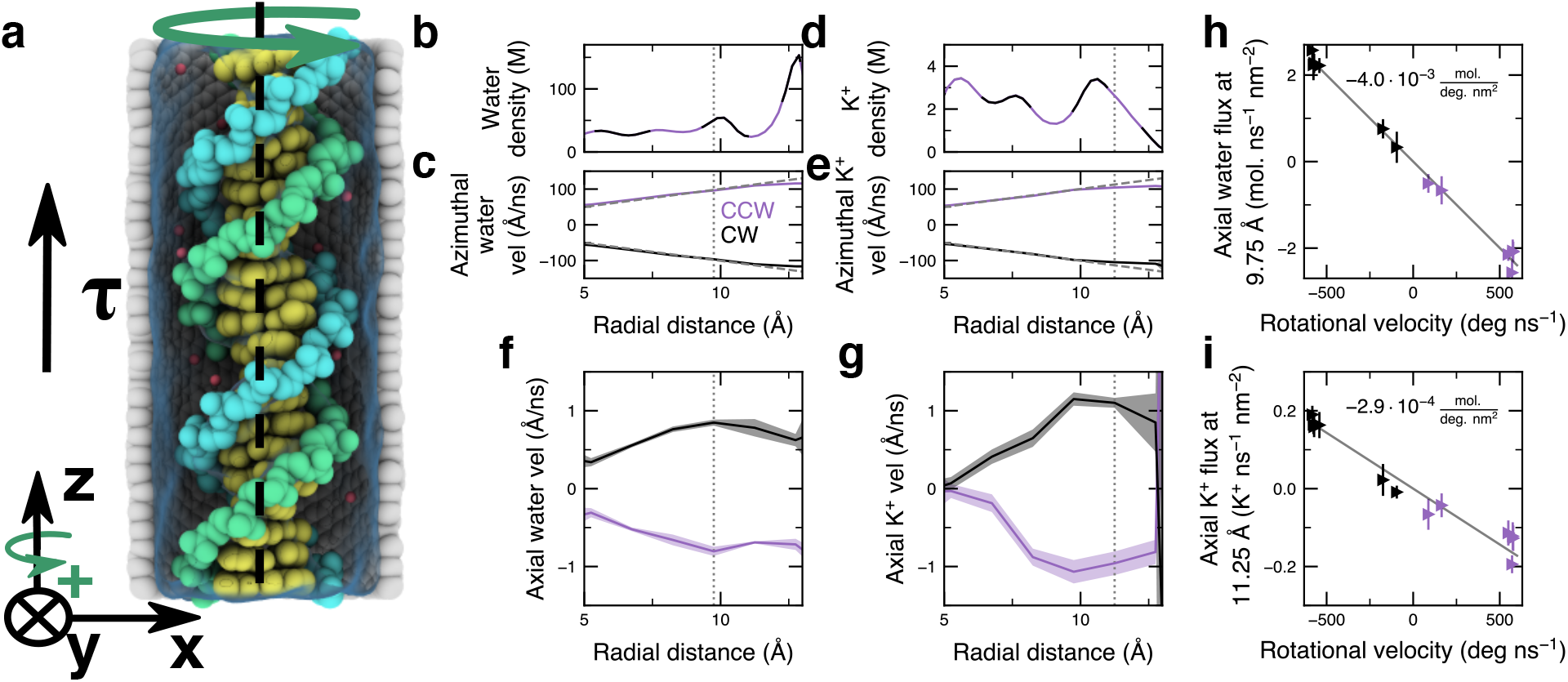
Flux generation mechanism. **a**, Cutaway view of the simulation system containing a 21 bp DNA helix within a 3.3 nm diameter carbon nanotube. Both nanopore and DNA are effectively infinite *via* bonds over the periodic boundary. Torque (*τ*) is applied along the nanopore axis (*z* axis) to the phosphorus atoms of the DNA to produce its rotation. **b**, Local density of water within the nanotube *versus* radial distance from the DNA axis. Here and elsewhere, the vertical dashed lines represent a distance of either 9.75 or 11.25 Å away from the DNA axis. In panels b–g, data were averaged over three MD trajectories, each at least 400 ns long, under a torque of *±*55.1 pN nm. The shaded regions represent the standard error of the mean computed between the three MD trajectories. CW and CCW and the associated colors denote the clockwise and counterclockwise rotation of the duplex. The bin size for radial averaging was 0.25 Å for data in panels b and d and 1.5 Å for data in panels c and e-i. **c**, Azimuthal water velocity *versus* radial distance from the DNA axis. **d**, Density of cations *versus* distance from the DNA axis. **e**, Azimuthal cation velocity *versus* distance from the DNA axis. **f**, Axial component of water velocity *versus* distance from the DNA axis. **g**, Axial component of cation velocity *versus* distance from the DNA axis. **h**, Axial water flux 9.75 Å away from the DNA axis *versus* rotational velocity. **i**, Axial flux of cations 11.25 Å away from the DNA axis *versus* rotational velocity. Each data point in panels h and i represents the average of a 400 ns MD trajectory. The error bars depict the standard error of mean calculated by splitting the trajectory in 50 ns blocks.

Driven by the rotation of the DNA duplex, water is seen to rotate about the DNA axis within the carbon nanotube, with the azimuthal velocity linearly increasing with the radial distance Fig. 2c. We attribute this behavior to nearly frictionless slipping of the adsorbed water along the carbon nanotube surface. Closer inspection, however, reveals the azimuthal velocity profile deviates slightly from ideal frictionless behavior (dashed lines in Fig. 2c) near the nanopore surface. Reflecting complexity of ion interactions with the grooves of the DNA, the average density of potassium ions exhibits non-trivial dependence on the radial distance, until falling off 11.25 Å away from the DNA axis, Fig. 2d. The azimuthal velocity of potassium ions, Fig. 2e, increases linearly with the radial distance, matching the azimuthal water velocity profiles.

Reflecting the conversion of a DNA duplex rotation into solvent flow through the nanopore, the axial component of water velocity is approximately 100 times smaller than the respective azimuthal component, Fig. 2f, and reaches a maximum about 9.75 Å away from the DNA axis. The axial velocity of potassium ions is also much smaller than their azimuthal velocity, Fig. 2g, and reaches a maximum proximal to the charged DNA backbone. Surprisingly, the axial velocity of potassium ions is approximately 1.5 times greater than that of water, Fig. 2f,g. The increased axial velocity of potassium compared to that of water suggests that the DNA duplex rotation applies a stronger force to the ions compared to water and that the water molecules follow the cations. The axial flux of water and ions are both found to scale linearly with the angular velocity of the DNA duplex rotation, Fig. 2h,i. Note that, although cations travel faster along the duplex axis than water, the higher number density of water results in higher axial flux, Fig. 2h,i. Qualitatively similar dependence of local water and ion velocity on radial distance was observed in MD simulations of an infinite DNA duplex rotating in bulk electrolyte solutions, Fig. S1.

To determine how electric charge at the nanopore surface can regulate the rotation-driven transport of water and ions, we built two variants of a previously described carbon nanopore system (Fig. 1a) by assigning partial charges to the atoms comprising the nanopore surface, Fig. 3a, see SI Methods for details. Charging the nanopore surface did not substantially alter the velocity of DNA duplex rotation under the same applied torque, Fig. S2a and Supporting Video S4 and 5, nor did it alter the average water flux driven by the rotation, Fig. 3b, despite slight increase in water adsorption at the positively charged nanopore surface, Fig. S3a. The charge on the nanopore surface was observed to produce the expected effect^58^ on ion concentration within the nanopore, with anion concentration increasing and cations concentration decreasing in the positively charged nanopore and *vice versa* in the negatively charged one, Fig. S3b,c. Consequently, the rate of rotation-driven transport of K^+^ ions increased in the negatively charged nanopore and decreased (but remained statistically significant) in the positively charged one in comparison to the neutral pore baseline, Fig. 3c. No significant rotation-driven transport of Cl^*−*^ ions was observed in the negatively charged or neutral nanopore systems, in contrast to the positively charged system where the transport was substantial, Fig. 3d. Note that the differential modulation of K^+^ and Cl^*−*^ transport by the pore charge originates from the presence of DNA that carries a substantial negative charge on its backbone.

**Figure 3:**
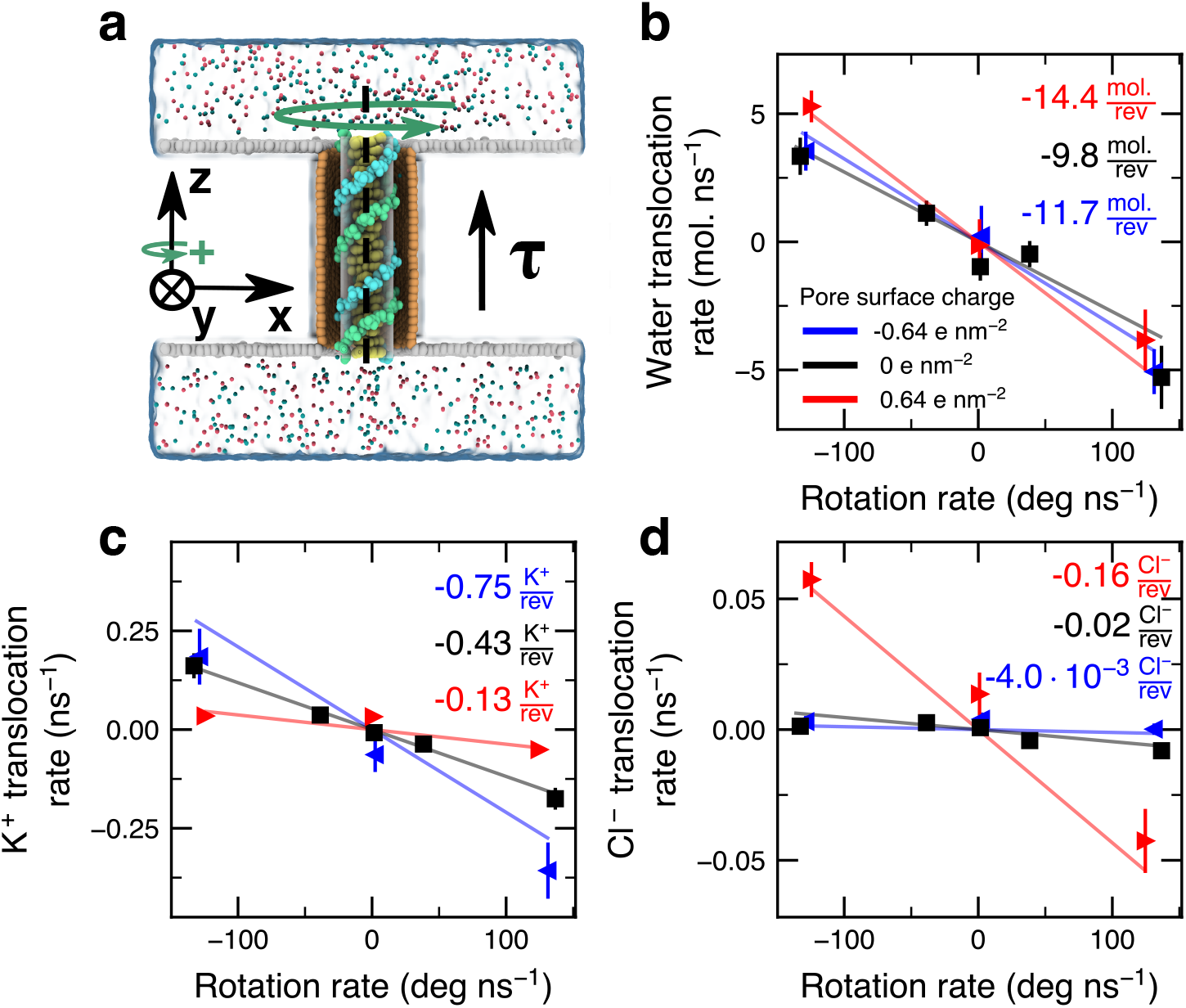
Effects of nanopore charge on torque-driven flux. **a**, Cutaway view of the simulation system, where a 21-bp DNA duplex is confined within a carbon nanopore and surrounded by 1 M KCl electrolyte. Atoms of the nanopore (shown in orange) are assigned partial charge to produce *±*0.64 e nm^*−*1^ surface charge density, where e is the charge of a proton. A torque *τ* is applied along the *z*-axis, rotating the DNA helix about its axis. **b–d**, Water (panel b), potassium (panel c) and chloride (panel d) transport rates as a function of DNA rotational velocity for charged and neutral nanopore systems. Each data point in panels b-d indicates the average value from a 500 ns molecular dynamics trajectory with the error bars representing the standard error of mean calculated from 50-ns block-averaged data.

Subsequent simulations of the neutral carbon nanopore systems containing 1 M of LiCl or NaCl revealed a modest effect of the monovalent cation type on the rotation-driven transport of water and ions, Fig. S4. The total number of water molecules as well as the radial water density, Fig. S5a, were nearly identical for all monovalent cation types, whereas the radial distribution of cations showed statistically significant differences, Fig. S5b. Thus, in agreement with a previous study,^59^ more Li^+^ and Na^+^ ions were observed to enter the DNA grooves in comparison to K^+^ likely because of their lower hydration.^60^

Repeating the DNA rotation simulations in 1 M MgCl_2_ or 1 M CaCl_2_ electrolyte revealed a modest decrease of the average water transport rate, Fig. 4a and Fig. S6, whereas the rotation-driven transport of divalent cations was reduced by 75–93% compared to that observed for monovalent electrolytes, Fig. 4b. Analysis showed that the presence of divalent cations does not affect the radial water density significantly, Fig. S5. We attributed the substantial reduction in the rotation-driven transport of divalent ions to their lower mobility and higher affinity for DNA,^59^ which draws the ions into close association where their axial velocity is negligible, Supporting Video S6.

**Figure 4:**
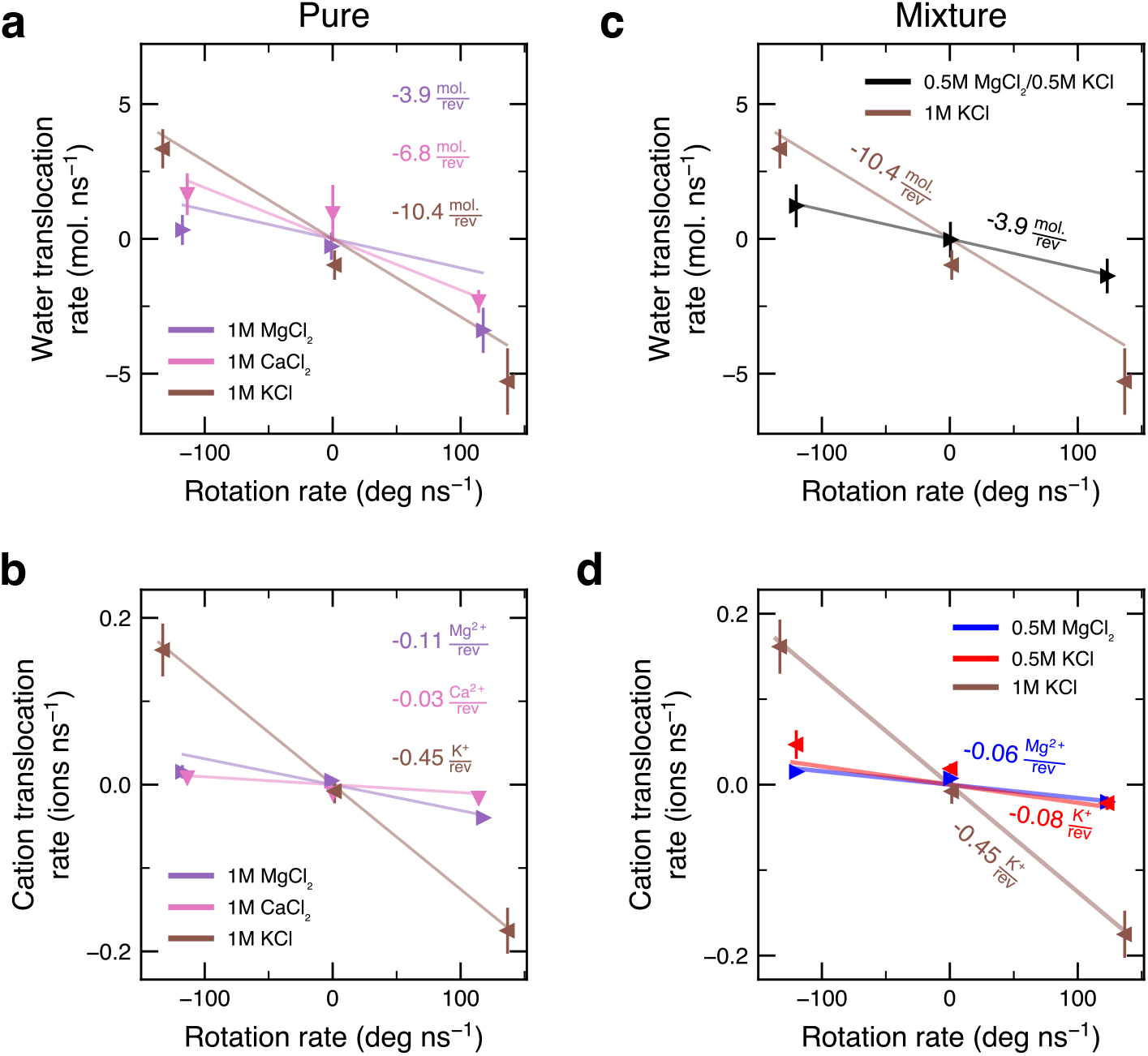
The effects of divalent cations on torque-driven flux. **a**,**b**, Water (panel a) and cation (panel b) transportation rates *versus* DNA rotational velocity in the systems solvated with divalent cations. **c**,**d**, Water (panel c) and cation (panel d) transportation rates *versus* DNA rotational velocity in systems containing a mixture of 0.5 M KCl and 0.5 M MgCl_2_. Each data point represents the average value from a molecular dynamics trajectory of at least 500 ns; the error bars are the standard error of the mean calculated from 50 ns block averaged data. For comparison, all panels also depict the transportation data for 1 M KCl solution.

Motivated by the observed dependence of the rotation-driven flux on the cation valence, we investigated if such an effect could be used for ion separation. Our simulation system contained our standard carbon nanopore solvated with a mixture of 0.5 M KCl and 0.5 M MgCl_2_ electrolytes, Fig. S7a and Supporting Video S7. The simulation of the mixed electrolyte system showed similar rates of the rotation-driven water flow as in pure divalent electrolyte systems, Fig. 4a,c, and nearly identical radial profiles of water density, Fig. S5a and Fig. S7c. However, the flux of K^+^ decreased 7-fold in comparison to that in a pure 1 M KCl system, while the flux of magnesium nearly halved compared to that in a pure 1 M MgCl_2_ system. While the reduction of magnesium flux is mostly accounted for by 2-fold reduction of Mg^2+^ density in the mixed system, Fig. S7d, the 7-fold reduction of K^+^ flux was unexpected. Thus, although the DNA duplex produces much higher flux of K^+^ than of Mg^2+^ in pure electrolyte systems, in the mixed electrolyte scenario, Mg^2+^ interactions with DNA and water suppress the DNA-rotation driven transport of K^+^.

To determine how electrolyte concentration affects the rotation-driven transport of ions and water, we built three additional systems containing 0.15, 0.5 and 2 M KCl electrolyte, Fig. S8. The transport efficiency of water and cations, defined here as the number of transported molecules per DNA duplex rotation, did not change statistically with bulk electrolyte concentration, whereas the transport efficiency of anions increased from being statistically near zero to a small, yet statistically significant, value at 2M KCl, Fig. S8. The apparent insensitivity of the rotation-driven transport to bulk electrolyte concentration is caused by the recruitment of counterions into the pore to neutralize the fixed DNA charge.

To investigate whether a rotating DNA duplex can indeed act as an Archimedes screw, we built a nanopore system, Fig. 5a, where an ion concentration gradient was maintained during the simulation *via* an external potential,^61^ see SI Methods for details. The steady state concentration of KCl electrolyte in these simulations was 0.6 and 1.2 M, respectively, below and above the membrane, Fig. 5b,c. The chemical potential created by the concentration difference between the two compartments (*∼*0.4 kcal mol^*−*1^) was seen to generate a nanopore ionic current of *−*27 pA K^+^ and 1.2 pA for Cl^*−*^, Fig. 5e,f and Supporting Table S1. Based on the results of our previous simulations (Fig. 1), pumping ions against the established concentration gradient would require the application of a negative torque.

**Figure 5:**
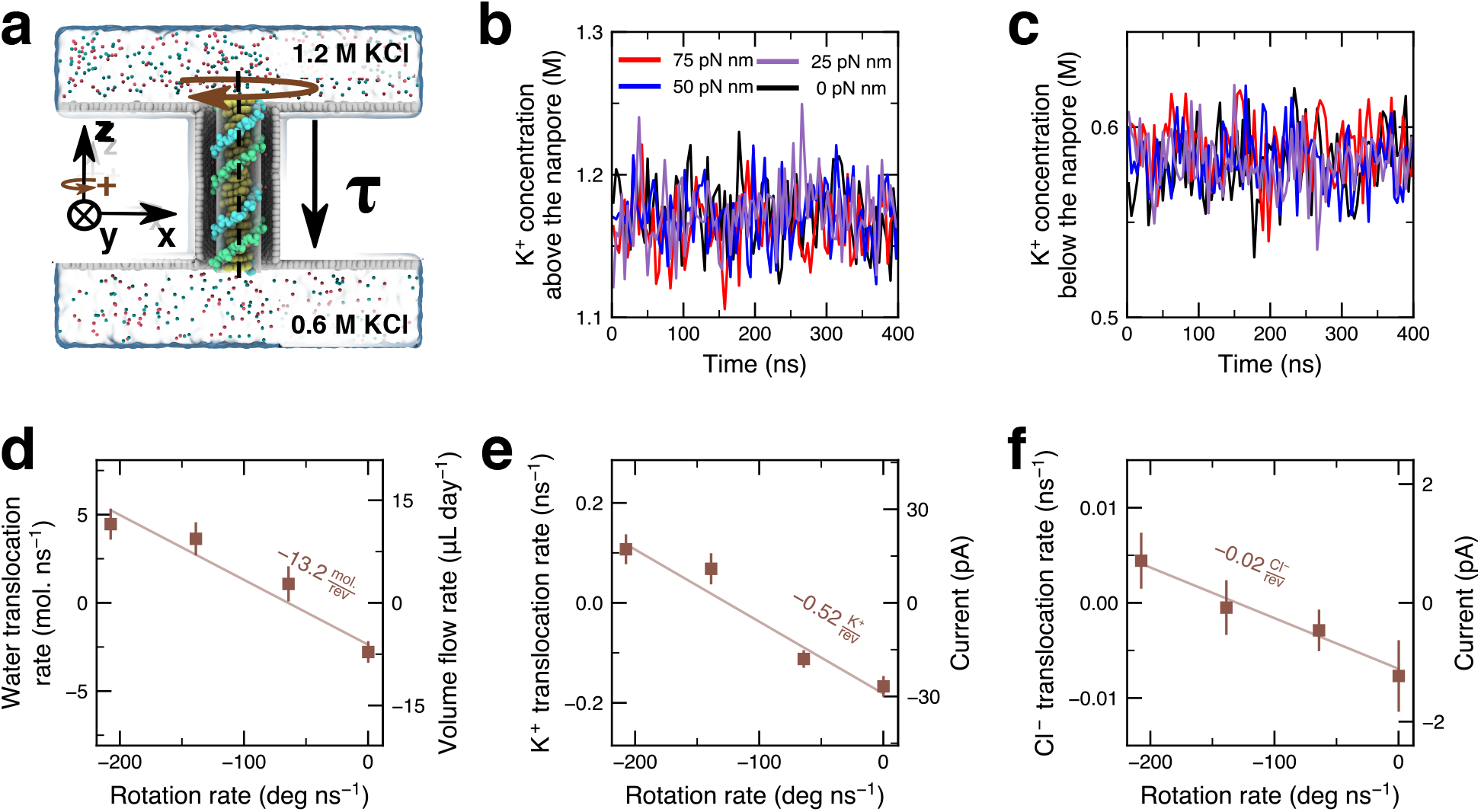
Pumping ions against concentration gradient. **a**, Cutaway view of the simulation system, containing a 21 bp DNA duplex held within a carbon nanotube 6.5 nm long and 3.3 nm in diameter. An external potential is applied to impose a gradient of ion concentration across the membrane (see SI Methods for details). **b**,**c**, Timeseries of the K^+^ density in the upper (b) and lower (c) compartments. The concentrations were averaged over the regions located 5Å away from the respective membrane surface and the periodic boundary along the *z* axis. The plots depict 4 ns block-averaged data. **d-f**, Rate of rotation-driven transport of water (d), K^+^ (e), and Cl^*−*^ (f) ions. A positive translocation rate describes a transport of molecular species from the bottom to the top compartment, *i*.*e*., against the concentration gradient. Each data point depicts the average value from an MD trajectory at least 400 ns. The error bars show the standard error of the mean calculated from 50 ns block averaged data.

Fig. 5d-f presents the obtained dependence of water, K^+^ and Cl^*−*^ ion translocation rate of the applied torque. In the absence of torque, water was found to flow from the top compartment to the bottom one, Fig. 5d, driven by the K^+^ selectivity of the ionic current, Fig. 5e,f. The emergence of such an electro-osmotic flow is well documented in the nanopore literature.^62,63^ As the magnitude of the torque increases, the water and ion flux reverses direction, attaining positive values at a DNA duplex rotation rate that exceeds 126 deg. ns^*−*1^, or when torques larger than *∼* 47 pN nm are applied to the DNA duplex. That is, the rotating DNA helix can indeed act as an Archimedes’ screw, moving water and ions against a chemical potential and concentration gradient.

In summary, we have shown through simulations how the application of torque to a DNA duplex can drive the flow of water and ions through a nanopore. Combined with our previous work that has shown generation of torque by a DNA duplex when subject to electric field or flow,^51^ our current work confirms reversibility of nanoscale molecular motors, in line with the previous studies of FoF1 ATP synthase and similar motors.^28^ Furthermore, we have shown that a realistic biological torque of *∼*50 pN nm can transform a DNA duplex confined within a nanopore into a molecular pump capable of transporting water and ions against a two-fold ion concentration gradient. While the efficiency of such an ion pump is considerably lower than that of biological molecular pumps, we envision the usage of helical molecular pumps built from DNA, RNA, PNA or other programmable chiral polymers to selectively bind and transport low abundance / high-value chemicals for separation, purification or concentration of biological matter.

## Acknowledgement

This work was supported by a grant from the National Science Foundation (ID-2411133). The supercomputer time was provided through the ACCESS allocation MCA05S028.

## Supporting Information Available

Simulations Methods detailing system setup, protocols and analysis; Supplementary Figures depicting DNA rotation and water and ion concentrations and fluxes under various conditions; Supplementary table detailing computed ionic currents and water fluxes; and Supplementary Videos depicting the simulations. This information is available free of charge via the Internet at http://pubs.acs.org.

