## Supplementary Information for "DNA: A Nanoscale Archimedes’ Screw"

### Supporting Information

Xavier Mleziva,[†,‡] Christopher Maffeo,[†,‡] and Aleksei Aksimentiev[*,†,‡]

†*Department of Physics; University of Illinois at Urbana–Champaign; Urbana, IL 61801, USA.*

‡*Beckman Institute for Advanced Science and Technology, University of Illinois at Urbana–Champaign; Urbana, IL 61801, USA.*

### Materials and Methods

**General MD methods.** All MD simulations were carried out using NAMD 2.14 program[1] under periodic boundary conditions. CHARMM36 parameters[2,3] were used to describe DNA and ions along with the CUFIX corrections[4] to DNA–ion interactions and the TIP3P model for water.[5] A particle-mesh Ewald scheme with a grid spacing of 1.2 Å was used to calculate the long-range electrostatic interactions.[6] Van der Waals and short-range electrostatic forces were evaluated using a 7–8 Å cutoff scheme. The simulations used the hydrogen mass repartitioning method,[7] which allowed us to set the integration time step to 4 fs. The full electrostatics were evaluated at each time step.[7] Energy minimization used a conjugate gradient method.[8]

**Assembly and equilibration of a DNA duplex in a carbon nanopore.** Carbon nanopore systems were built using the InorganicBuilder plugin of VMD[9] as described previously.[10] The nanopore was constructed by duplicating a single-layer hexagonal graphene sheet, then shifting one of them by 6.5 nm relative to the other, normal to the planes of the sheets. A hexagonal pore with a circumscribed radius of 1.6 nm was cut into the center of each sheet. The atoms along the edge of each pore were then connected to the ends of an achiral carbon nanotube of a matching diameter and 6.5 nm in length. Building nanopores this way introduced defects at each nanotube/graphene sheet junction.[10] Charged nanopore systems were produced by assigning a partial charge of $\pm 0.0163\ e$ to each carbon atom of the nanotube, which corresponds to a surface charge density of $\pm 0.64\ e\,\mathrm{nm}^{-2}$

A 21 bp-long DNA duplex was constructed using mrDNA[11] to have the d(ATCG)$_5$dA sequence. The duplex was placed coaxially inside the nanopore. The `Solvate` and `Autoionize` plugins of VMD were used to add water and ions, producing a range of systems differing by the composition and concentration of the electrolyte solution. No solvent molecules were placed in the toroidal volume surrounding the nanopore between the two graphene sheets.

During all simulations, all nanopore atoms (including the two carbon sheets) were harmonically restrained to their initial coordinates ($k_{\mathrm{spring}}$ = 1.0 kcal mol$^{-1}$ Å$^{-2}$), except for the atoms at the upper nanotube/graphene junction. Harmonic bond restraints ($k_{\mathrm{spring}}$ = 1.0 kcal mol$^{-1}$ Å$^{-2}$; 2.9 Å rest length) were placed between the purine N1 and pyrimidine N3 atoms of each basepair to prevent the possibility of DNA duplex melting. Each system was minimized for 2,400 steps while harmonically restraining ($k_{\mathrm{spring}}$ = 1.0 kcal mol$^{-1}$ Å$^{-2}$) the non-hydrogen atoms of the duplex about their initial coordinates. Following the minimization, the systems were equilibrated for 14.4 ns under the harmonic constraints in the constant number of atoms, pressure and temperature (NPT) ensemble using the Nosé-Hoover Langvin piston, with pressure and temperature targets of 295 K and 1 atm. The Langevin piston had a decay rate of 2000 and a period of 1000. Each system's dimension along the DNA axis ($z$-axis) was allowed to fluctuate during equilibration to achieve the pressure target, while the orthogonal dimensions were fixed. The Langevin damping coefficient was 0.1 ps$^{-1}$.

**Assembly and equilibration of bulk-solution periodic DNA duplex systems.** An all-atom model of a d(ACTG)$_5$dA duplex DNA was constructed using the nucleic acid builder.[12] The duplex was submerged in a 64×64×70 Å$^3$ volume of 1 M KCl electrolyte. The DNA strands were connected to themselves over the periodic boundary using the LKNA patch. The systems were minimized and equilibrated as described above except that only the phosphorus atoms in the duplex were harmonically restrained ($k_{\mathrm{spring}}$ = 1.0 kcal mol$^{-1}$ Å$^{-2}$) to their initial coordinates and no additional restrains were used to enforce the base pairing. Upon assembly and energy minimization, the bulk solution systems were first equilibrated in the constant number of atoms, volume, and temperature (NVT) ensemble for 14.4 ns before a 14.4 ns equilibration in the NPT ensemble. During the NPT equilibration, the systems were allowed to change their dimensions along all three Cartesian axes, in a way that preserved the ratio between all three axes. The Langevin piston had a decay rate of 100 and a period of 50.

**Assembly of a periodic DNA duplex in an infinite nanotube.** An all-atom model of an achiral carbon nanotube 3.3 nm in diameter and 7 nm in length was built using the InorganicBuilder plugin of VMD.[9] A custom script connected the carbon atoms of the nanotube over the periodic boundary of the system. The periodic model of a 21-bp DNA duplex (described above) was placed coaxially inside the nanotube. Water and ions were added to the nanotube/DNA system matching the water and ion densities observed during a 14.4 ns equilibration of the same-diameter nanotube exposed to 1 M KCl bulk electrolyte. Specifically, 43 potassium ions and one chloride ion were added to produce an electrically neutral system. The resulting system (main text Fig. 2a) was minimized for 2,400 steps and equilibrated in the constant number of particle, volume and temperature (NVT) ensemble for 14.4 ns before the NPT equilibration. During the NPT equilibration, no harmonic restraints were applied to atoms of DNA and the nanotube and the system was allowed to change its dimensions along the $z$-axis, preserving the cross-sectional area of the $x$–$y$ plane. The Langevin piston had a decay rate of 100 and a period of 50. After the NPT equilibration, the system's coordinates were adjusted to center the nanotube at the origin and align it coaxially with the $z$-axis. The target coordinates of the harmonic restraints were adjusted to accommodate a slight ($\sim$0.06%) compression of the nanotube observed during the equilibration.

**Applied torque simulations.** All applied torque simulations were conducted in the NVT ensemble. The system's dimensions were set to the NPT equilibration average. The same `tclForces` script harmonically restrained and imparted torque on each phosphorus atom of the DNA molecule. One set of harmonic restraints ($k_{\mathrm{spring}}$ = 1.0 kcal mol$^{-1}$ Å$^{-2}$) confined the motion of the phosphorus atoms to the surface of a 9.4 Å-radius cylinder. Another set of harmonic restraints (same spring constants) confined displacement of the phosphorus atoms along the $z$-axis. Taken together, these restraints kept the DNA molecule centered in the pore, while allowing the molecule to rotate about the $z$ axis. In nanopore simulations, additional harmonic restraints (between the N1 atoms of purines and the N3 atoms of pyrimidines) enforced base pairing within the DNA duplex ($k_{\mathrm{spring}}$ = 1 kcal mol$^{-1}$ Å$^{-2}$; 2.9 Å rest length). For a given value of torque, $\tau_{\mathrm{tot}}$, the `tclForces` script applied the following force to each phosphorus atom of the duplex: $F_{\tau} = \tau_{\mathrm{tot}}/(r \cdot N_{\mathrm{p}})$, where $r$ is the radial distance of the atom ($\sim$9.4 Å) and $N_{\mathrm{p}}$ is the number of phosphorus atoms in the DNA duplex. The coordinate frames were recorded every 2400 steps, while the torque and restoring forces were recalculated every 200 steps except for the infinite nanotube system, where the restoring forces were recalculated every 100 steps.

**Simulation under ion concentration gradient.** The 2:1 ratio of electrolyte concentration was realized following a method introduced earlier.[13] The `gridforces` function of NAMD[14] was used to apply forces to direct ions from the lower reservoir into the upper reservoir (main text Fig. 5a). The force magnitude was determined from the chemical potential difference of the two reservoirs

$$\mu = -kT \ln\left(\frac{C_{\text{low}}}{C_{\text{high}}}\right),$$

where $C_{\text{low}}$ and $C_{\text{high}}$ are the target concentrations of the solution. For our conditions, the calculation yields $\mu = 0.406$ kcal mol$^{-1}$. A custom script was then utilized to produce a grid potential with a spacing of 0.5 Å and a height of 7 Å, featuring a 5 Å region where the potential ramps from zero to the maximum value (0.406 kcal mol$^{-1}$). The grid potential was placed in the upper reservoir such that the forced region was 2 Å from the periodic boundary of the system. The simulation systems were built and equilibrated according to the nanopore construction protocol described above, except that the upper reservoir was solvated with 1 M KCl, while the rest of the system was solvated with 0.5 M KCl. Prior to production simulations, the systems (main text Fig. 5a) were equilibrated in the NPT ensemble for 1.44 ns in the absence of the external potential, followed by another 14.4 ns NPT equilibration under the gridforces potential, which was sufficient to establish the desired concentration gradient.

**Data Analysis.** The transport of water and ions was characterized using a previously described method.[15] Briefly, the instantaneous current of the target molecular species was calculated as

$$I(t) = \frac{1}{Ldt}\sum_{i=1}^{N} q_i dz_i(t),$$

where $L$ is the length of the sampled region along the $z$ axis, and $dz_{\text{i}}$ and $q_{\text{i}}$ are the $z$-displacement and charge of molecule $i$, respectively, and the sum runs over all molecules of interest in the sampled region. Note that $dz_{\text{i}}$ is computed as a finite difference between frames and only includes the displacement that intersects the region of interest, *i.e.*, the nanopore volume at least 15 Å away from either nanopore entrance. The instantaneous transport of water was characterized by analyzing the motion of water oxygen atoms and setting their charge to unity.

For systems containing periodic DNA, we used a previously described method[10,16] to calculate the velocity of molecular species in a spatially resolved manner. For a given molecular species, a three-frame, centered finite difference displacement vector was computed for each atom in the cylindrical coordinate systems ($\hat{r}$, $\hat{\theta}$, and $\hat{z}$) centered on the DNA. The displacement vectors were assigned to 1.5 Å radial bins according to the atom's radial coordinate in the central frame. The density of atoms in bin $i$ was calculated as

$$\rho_i = \frac{n_i}{2\pi r_i \Delta r h},$$

where $r_i$ is the radius of the center of the bin, Δr is the bin's width (1.5 Å), $h$ is the height of the measurement region and $n_i$ is the number of atoms in bin $i$. The bulk reservoir concentrations were calculated by dividing the number of molecules in a given region by the volume of that region. For a given bin, the average velocities were computed by dividing the sum of all atoms' displacements by the time interval between the first and last frames and by the number of atoms in that bin. The radius-dependent flux was computed by dividing the sum of all atoms' velocities in a bin by the volume of that bin. The rotational velocity of DNA was calculated by first computing the frame-to-frame angular displacement about the $z$ axis of a phosphorus atom at the end of the duplex. The angular displacement for each frame was set to have a value between $-\pi$ and $\pi$. The cumulative angular displacement was then calculated as a sum of the adjusted angular displacements.

All linear fits were obtained using a standard linear regression assuming a $y$-intercept at the origin, except for the simulations under a concentration gradient (main text Fig. 5d-f), where the intercept was treated as a fitting parameter.

### Supporting Figures and Table

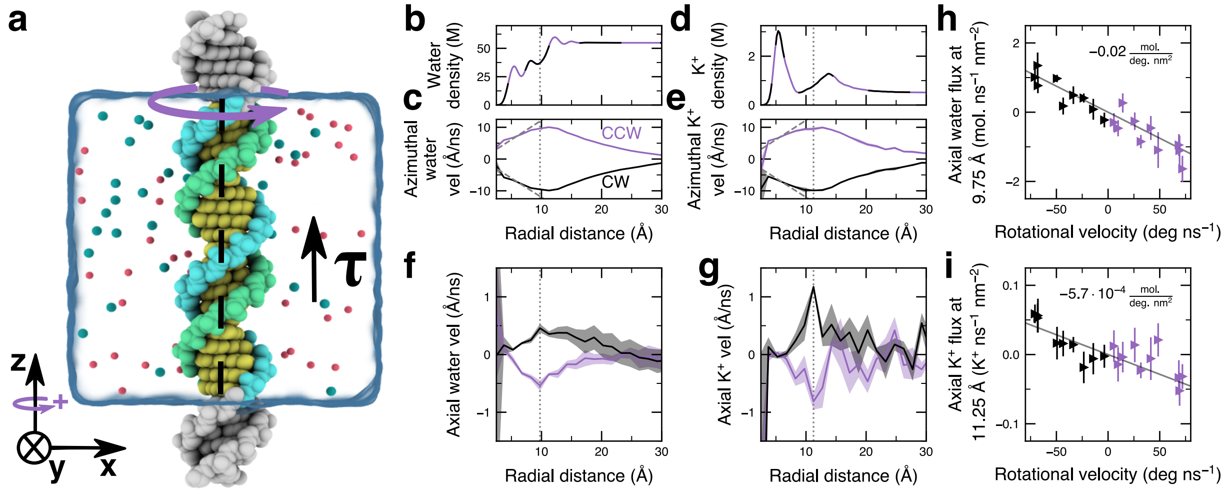


**Figure S1: Mechanism of flux generation by DNA rotation in bulk solvent. a,** Simulation system containing a 21 bp DNA helix submerged in a 1 M KCl electrolyte. The DNA is effectively infinite *via* bonds over the periodic boundary. Torque ($\tau$) is applied along the DNA axis ($z$ axis) to the DNA phosphorus atoms to produce its rotation. **b,** Density of water *versus* radial distance from the DNA axis. Here and elsewhere, the vertical dashed lines represent a distance of either 9.75 or 11.25 Å away from the DNA axis. In panels b–g, data were averaged over three MD trajectories, each at least 400 ns long, under a torque of ±55.1 pN nm. The shaded regions represent the standard error of the mean computed between the three MD trajectories. CW and CCW and the associated colors denote the clockwise and counterclockwise rotation of the duplex. The bin size for radial averaging was 0.25 Å for data in panels b and d and 1.5 Å for data in panels c and e-i. **c,** Azimuthal water velocity *versus* radial distance from the DNA axis. **d,** Density of cations *versus* distance from the DNA axis. **e,** Azimuthal cation velocity *versus* distance from the DNA axis. **f,** Axial component of water velocity *versus* distance from the DNA axis. **g,** Axial component of cation velocity *versus* distance from the DNA axis. **h,** Axial water flux 9.75 Å away from the DNA axis *versus* rotational velocity. **i,** Axial flux of cations 11.25 Å away from the DNA axis *versus* rotational velocity. Each data point in panels h and i represents the average of a 400 ns MD trajectory. The error bars depict the standard error of mean calculated by splitting the trajectory in 50 ns blocks.

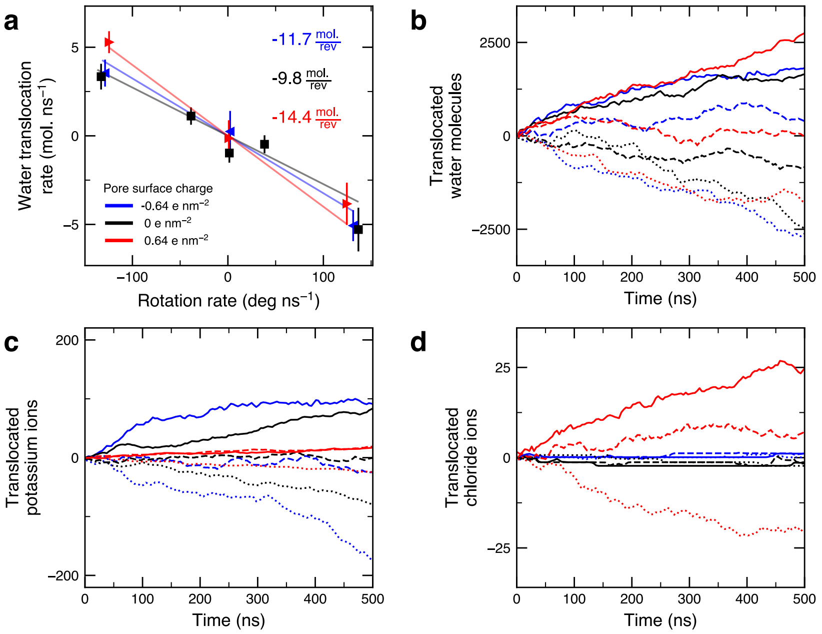


**Figure S2: Effects of nanopore charge on torque-driven rotation and flux. a,** Water translocation rate *versus* DNA rotation rate for carbon nanopore systems having a neural or positively or negatively charged surface. The systems is shown in main text Fig. 3a. Each data point represents the average of a 500 ns MD trajectory with the error bars representing standard error of mean calculated from 50-ns block-averaged data. **b-d,** Number of water molecules (panel b), potassium ions (panel c) and chloride ions (panel d) transported during MD simulations performed at the following values of torque: +50 (solid); 0 (dashed); or −50 (dotted) pN nm, applied to the DNA duplex threading the nanopore. The time series data were filtered using 4-ns block averaging.

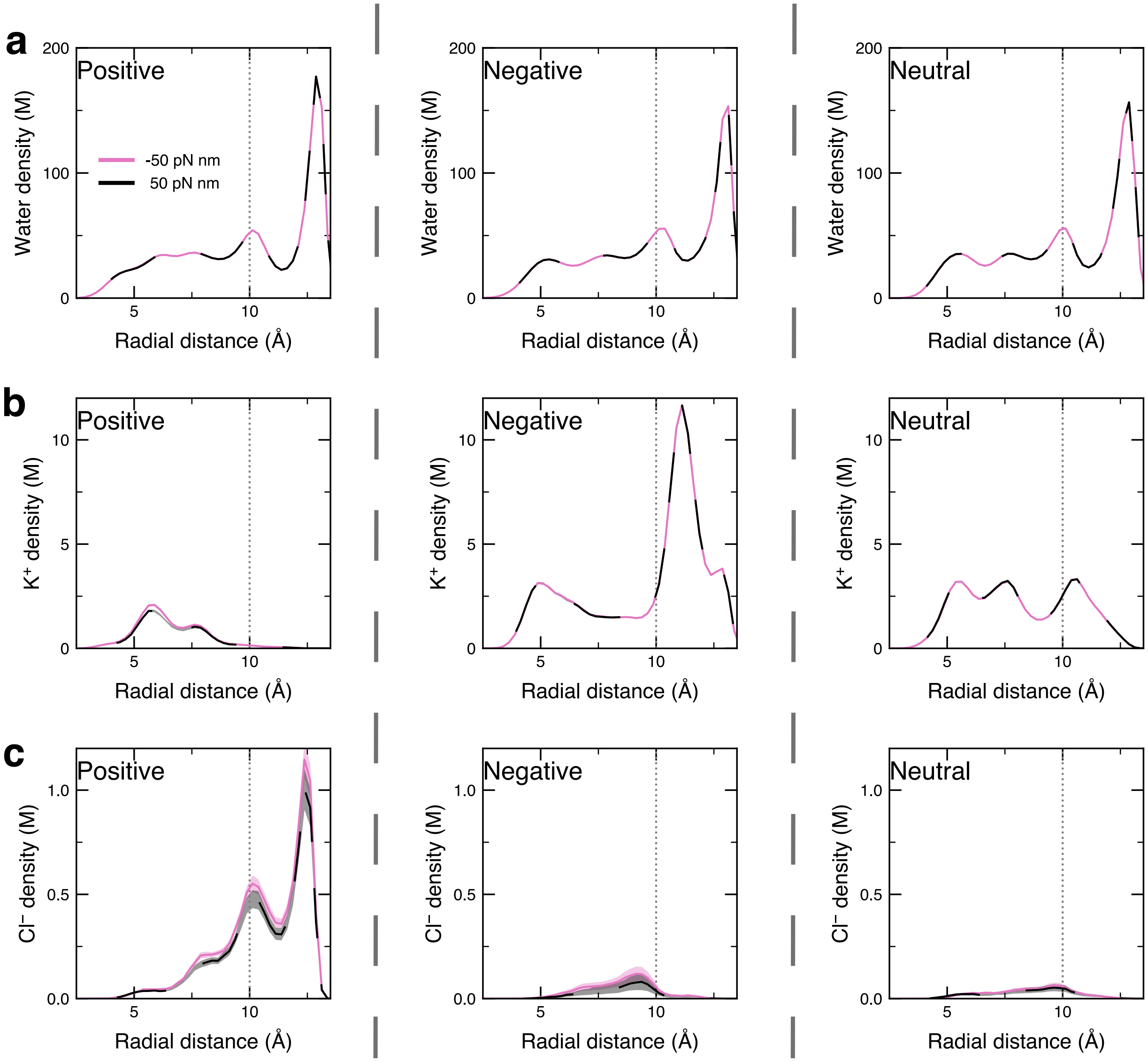


**Figure S3: Effects of nanopore charge on radial distribution of water and ions. a-c,** Average density of water (panel a), potassium (panel b), and chloride (panel c) ions within the nanopore *versus* distance from the DNA axis. Data are shown for MD simulations under ±50 pN nm torque applied to the DNA duplex. The density profiles were computed using 0.25 Å radial bins. The shaded regions represent the standard error of the mean computed from the respective MD trajectories, each being at least 500 ns. Positive (left), negative (middle), and neutral (right) labels corresponds to the nanopore surface charge density of +0.64, −0.64 and 0 e $nm^{-1}$.

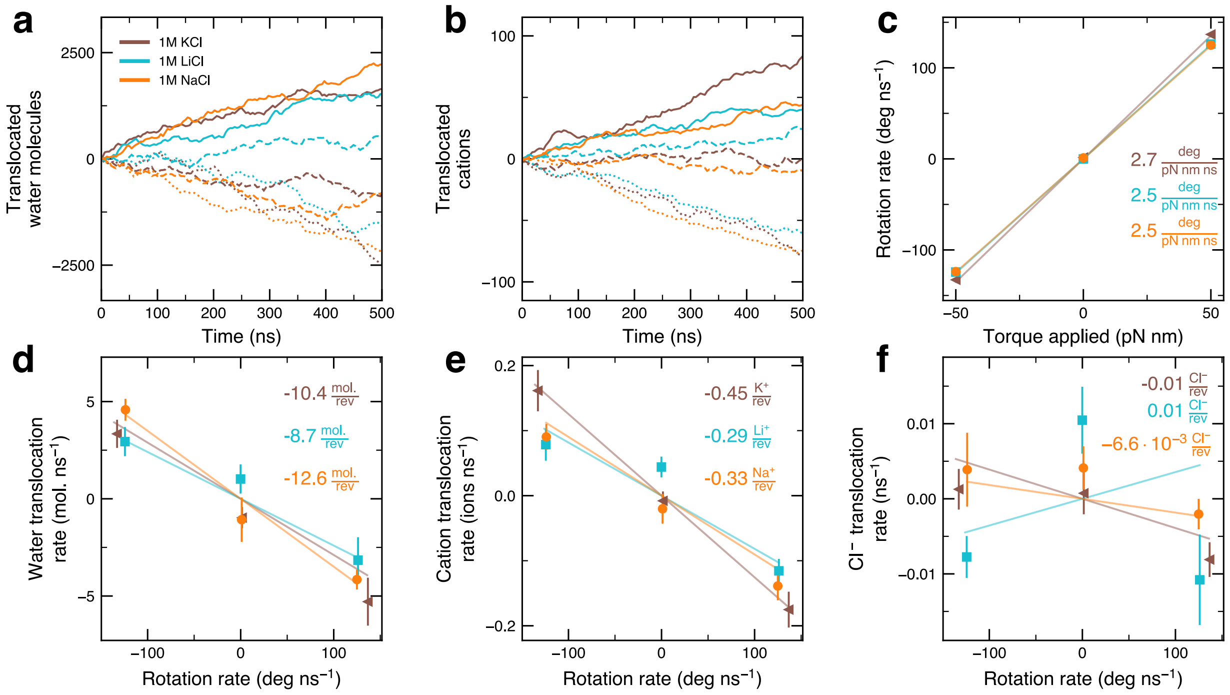


**Figure S4: Torque-driven flux in monovalent electrolytes. a,b,** Number of water molecules (panel a) and cations (panel b) transported through a nanopore during MD simulations performed at the following values of torque: +50 (solid); 0 (dashed); or −50 (dotted) pN nm, applied to the DNA duplex threading the nanopore. The time series data were filtered using 4-ns block averaging. **c,** DNA duplex rotation rate *versus* applied torque for simulations carried out in LiCl, NaCl, and KCl, the colors are defined in panel a. **d-f,** Water (panel d), cation (panel e) and chloride ion (panel f) transport rates as a function of DNA rotational velocity. In panels c-f, each data point represents an average over an MD trajectory of at least 500 ns. The error bars represent the standard error of the mean calculated by splitting the data into 50 ns blocks.

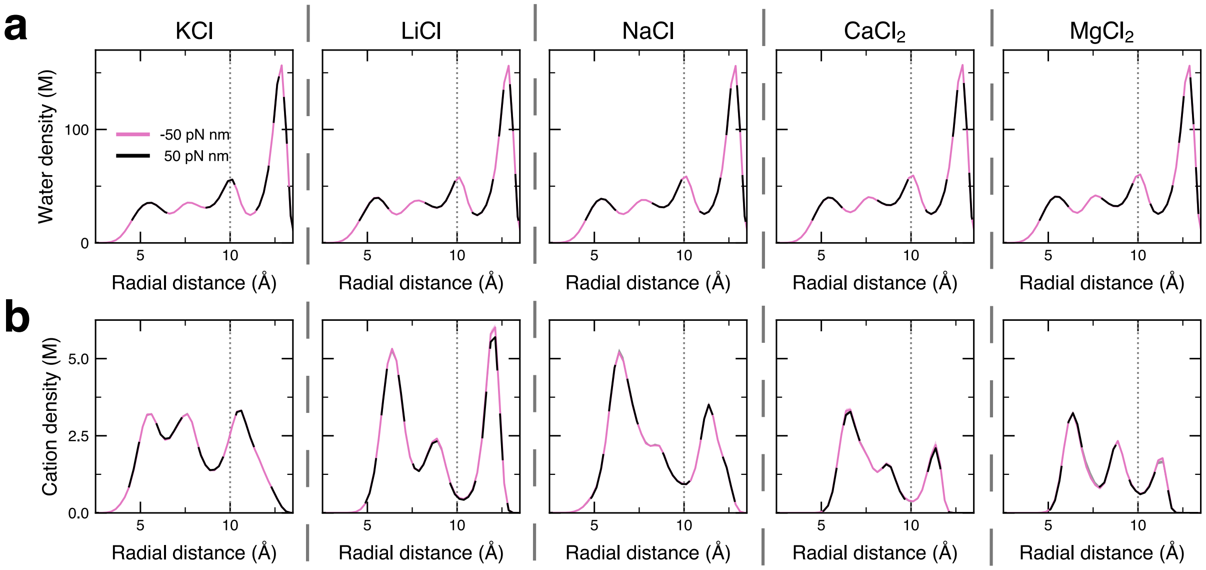


**Figure S5: Effects of cation type on radial distribution of water and cations. a,b**, Radial distribution of water (panel a) and cations (panel b) within the nanopore *versus* distance from the DNA axis. Each column depicts distributions with 0.25 Å bins averaged over MD simulations of at least 500 ns conducted under a torque of ±50 pN nm applied to the DNA duplex. The bulk ion concentration is 1 M in each system. Each column shows date for KCl, LiCl, NaCl, $CaCl_2$, and $MgCl_2$ electrolyte. At every condition, the standard error of the mean among the distributions computed after splitting the respective trajectory into 50-ns blocks is smaller than the line width.

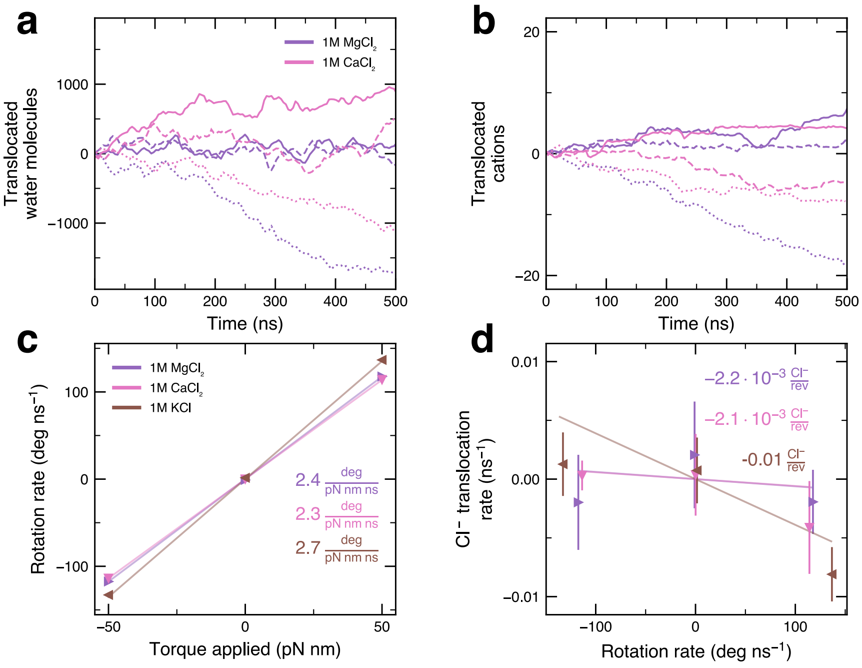


**Figure S6: Effects of divalent cation species on torque-driven flux. a,b,** Number of water molecules (panel a) and cations (panel b) transported through a nanopore during MD simulations performed at the following values of torque: +50 (solid); 0 (dashed); or −50 (dotted) pN nm, applied to the DNA duplex threading the nanopore. The time series data were filtered using 4-ns block averaging. **c,** DNA duplex rotation rate *versus* applied torque. **d,** Chloride ion transport rate as a function of DNA rotational velocity. In panels c and d, each data point represents an average over and MD trajectory of at least 500 ns. The error bars represent the standard error of the mean calculated after splitting data into 50 ns blocks.

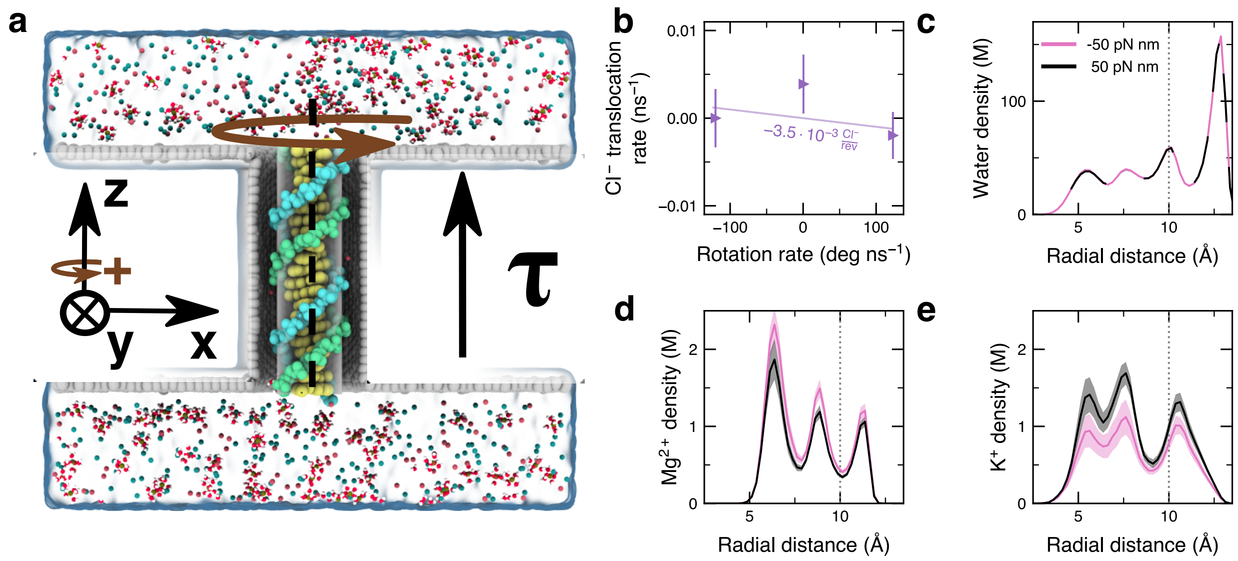


**Figure S7: Torque-driven flux in a mixed electrolyte solution. a,** Cutaway view of the simulation system, containing a 21 bp DNA duplex held within a carbon nanotube 6.5 nm long and 3.3 nm in diameter, submerged in an electrolyte containing 0.5 M KCl and 0.5 M $MgCl_2$. **b,** Chloride ion transportation rate *versus* rotational velocity. Each data point depicts the average value from the MD trajectory; the error bars depict the standard error of the mean calculated from 50-ns block averaged data. **c-e,** Radial description of water (panel c), potassium (panel d) and magnesium (panel e) ions inside the nanopore. Each distribution was computed using 0.25 Å radial bins by averaging over MD trajectories of at least 500 ns under ±50 pN nm torque applied to the DNA duplex. Shaded regions depict the standard error of mean among distributions computed after splitting the respective MD trajectory into 50-ns blocks.

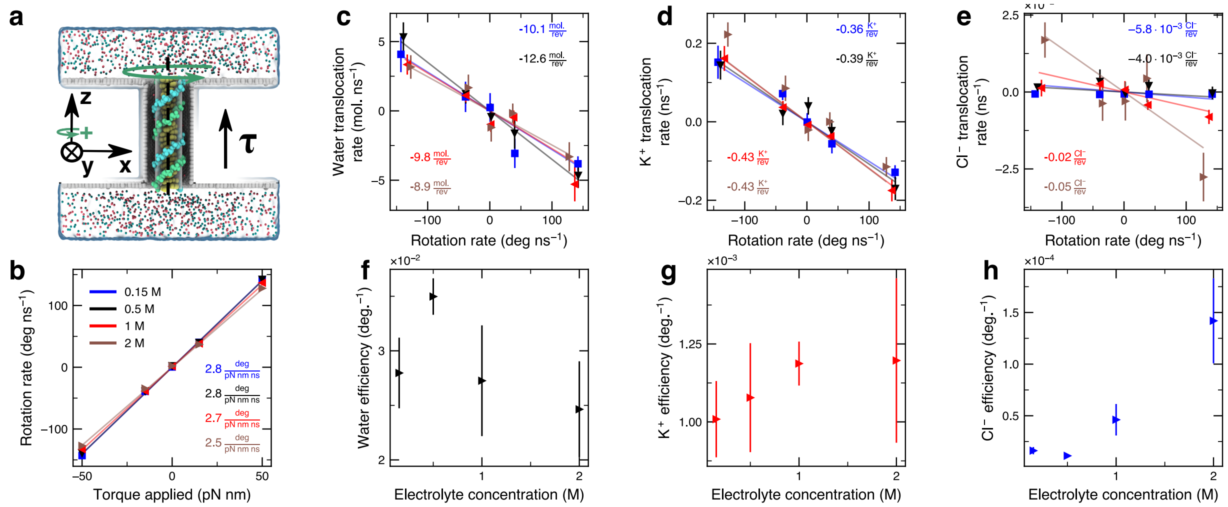

**Figure S8: Effects of KCl concentration on torque-driven flux. a,** Cutaway view of the simulation system, containing a 21 bp DNA duplex held within a carbon nanotube 6.5 nm long and 3.3 nm in diameter, submerged in 2 M KCl electrolyte. **b,** Rotational velocity of the DNA duplex *versus* torque applied to the duplex. **c-e,** Water (panel c), potassium (panel d) and chloride (panel e) transportation rates *versus* the DNA rotational velocity. In panels c-e, each data point represents the average value from a molecular dynamics trajectory of least 400 ns; the error bars represent the standard error of the mean calculated using 50-ns block averaged data. **f-h,** Transport efficiency of water molecules (panel f), potassium (panel g) and chloride (panel h) ions as a function of the bulk KCl concentration. Each data point in panels f-h depicts the efficiency value obtained as a linear regression fit to the dependence of the respective translocation rate on the rate of DNA rotation (panels c-e); the error bars indicate the standard error.

### Supporting Videos

**Supporting Video 1:** Rotation of a DNA duplex within an uncharged nanopore driven by a 50 pN nm torque applied to the DNA along its axis. The 21 bp duplex is shown using van der Waals (vdW) spheres (yellow bases; red and orange backbone). The ions within the 1 M KCl electrolyte are shown with vdW spheres in red ($K^+$) and blue ($Cl^-$). Phosphorus atoms of the duplex are harmonically restrained to the surface of a cylinder such that the DNA rotates in place about its central axis without drifting. The video depicts an 86 ns excerpt of a 500 ns molecular dynamics trajectory.

**Supporting Video 2:** Stochastic rotation of a DNA duplex within an uncharged nanopore in the absence of an applied torque. The 21 bp duplex is shown using vdW spheres (yellow bases; red and orange backbone). The ions within the 1 M KCl electrolyte are shown with vdW spheres in red

($K^+$) and blue ($Cl^-$). Phosphorus atoms of the duplex are harmonically restrained to the surface of a cylinder such that the DNA rotates in place about its central axis without drifting. The video depicts an 86 ns excerpt of a 500 ns molecular dynamics trajectory.

**Supporting Video 3:** Rotation of a DNA duplex within an uncharged nanopore driven by a $-50$ pN nm torque applied to the DNA along its axis. The 21 bp duplex is shown using vdW spheres (yellow bases; red and orange backbone). The ions within the 1 M KCl electrolyte are shown with vdW spheres in red ($K^+$) and blue ($Cl^-$). Phosphorus atoms of the duplex are harmonically restrained to the surface of a cylinder such that the DNA rotates in place about its central axis without drifting. The video depicts an 86 ns excerpt of a 500 ns molecular dynamics trajectory.

**Supporting Video 4:** Rotation of a DNA duplex within a positively charged nanopore driven by a 50 pN nm torque applied to the DNA along its axis. The 21 bp duplex is shown using vdW spheres (yellow bases; red and orange backbone). The ions within the 1 M KCl electrolyte are shown with vdW spheres in red ($K^+$) and blue ($Cl^-$). Phosphorus atoms of the duplex are harmonically restrained to the surface of a cylinder such that the DNA rotates in place about its central axis without drifting. The video depicts an 86 ns excerpt of a 500 ns molecular dynamics trajectory.

**Supporting Video 5:** Rotation of a DNA duplex within a negatively charged nanopore driven by a 50 pN nm torque applied to the DNA along its axis. The 21 bp duplex is shown using vdW spheres (yellow bases; red and orange backbone). The ions within the 1 M KCl electrolyte are shown with vdW spheres in red ($K^+$) and blue ($Cl^-$). Phosphorus atoms of the duplex are harmonically restrained to the surface of a cylinder such that the DNA rotates in place about its central axis without drifting. The video depicts an 86 ns excerpt of a 500 ns molecular dynamics trajectory.

**Supporting Video 6:** Rotation of a DNA duplex within an uncharged nanopore driven by a 50 pN nm torque applied to the DNA along its axis. The 21 bp duplex is shown using vdW spheres (yellow bases; red and orange backbone). The ions within the 1 M $MgCl_2$ electrolyte are shown with vdW spheres in dark blue ($Mg^{2+}$) and blue ($Cl^-$). Water molecules in magnesium hexahydrates are shown with vdW (red oxygen; white hydrogen). Phosphorus atoms of the duplex are harmonically restrained to the surface of a cylinder such that the DNA rotates in place about its central axis without drifting. The video depicts an 86 ns excerpt of a 500 ns molecular dynamics trajectory.

**Supporting Video 7:** Rotation of a DNA duplex within an uncharged nanopore driven by a 50 pN nm torque applied to the DNA along its axis and subject to an concentration gradient across the pore. The upper and lower volumes are maintained at 1.2 M and 0.6 M KCl electrolyte, respectively. The 21 bp duplex is shown using vdW spheres (yellow bases; red and orange backbone). The ions within the 1 M KCl electrolyte are shown with vdW spheres in red ($K^+$) and blue ($Cl^-$). Phosphorus atoms of the duplex are harmonically restrained to the surface of a cylinder such that the DNA rotates in place about its central axis without drifting. The video depicts an 86 ns excerpt of a 500 ns molecular dynamics trajectory.
